# Rapid CE–MS with Real-Time Eco–AI Resolves Proteomic Heterogeneity Among Single Human Neutrophils

**DOI:** 10.1101/2025.08.14.670204

**Authors:** Bowen Shen, Fei Zhou, Isabelle Luz, Laura G. Rodriguez, Steven J. Prior, Wagner Fontes, Peter Nemes

## Abstract

Single-cell proteomics by mass spectrometry is advancing rapidly, yet throughput and sensitivity remain limiting—particularly for small, protein-poor cell types such as neutrophils. As the most abundant circulating leukocytes in humans, neutrophils are central to immune defense and inflammation, but their proteomes comprehensive single-cell level characterization has only concurrently been reported^1^ and remains limited. Here, we introduce a rapid capillary electrophoresis–mass spectrometry (CE–MS) platform, which integrates electrophoresis-correlative real-time data acquisition with sub-7-minute separations and artificial intelligence (AI)-based data processing software to achieve deep, high-throughput profiling. Using single-cell–equivalent HeLa digests, the Rapid Eco–AI platform identified ∼1,350 proteins from 300 pg and ∼835 proteins from 75 pg of input— approaching the complexity of a mammalian cell proteome. Applied to freshly isolated human neutrophils, the workflow identified 151 proteins from ∼2 pg of material, ∼3% of the total cell proteome. Analysis of 13 individual cells revealed marked functional heterogeneity across pathways of degranulation, neutrophil extracellular trap (NET) formation, chemotaxis, and innate immunity, with hierarchical clustering resolving at least four distinct proteomic subtypes. These results establish Rapid Eco–AI as a sensitive, scalable, and broadly applicable CE–MS approach for immune-cell phenotyping at single-cell and subcellular resolution, facilitating new research opportunities in systems immunology and clinical proteomics.

## INTRODUCTION

Mass spectrometry (MS)–based proteomics has recently reached single-cell resolution, offering molecular snapshots that complement transcriptomics and refine our understanding of cellular states. However, most single-cell mass spectrometry (MS) workflows, the modern technology of choice for proteomics, still face a depth–throughput trade-off: barcoding and multiplexing accelerate sample preparation, yet long separations and slow tandem MS (MS²) duty cycles limit the number of cells that can be profiled in a day.

Neutrophils present a demanding test for any single-cell proteomics platform. These short-lived innate immune cells dominate the human peripheral leukocyte pool and play central roles in inflammation, host defense, and disease progression. Emerging evidence points to substantial functional and phenotypic heterogeneity, particularly in cancer, chronic inflammation, and immune dysregulation.^2–3^ Single-cell transcriptomics of >25,000 neutrophils has mapped this diversity,^4^ but corresponding protein-level data remain scarce, only one study so far^1^. Neutrophils are small (12–15 µm diameter) and protein-poor (∼60 pg total protein per cell),^5^ making their proteomic investigations analytically challenging. Early gel–nanoLC–MS studies required >10 million neutrophils to identify ∼250 proteins,^6^ and even the most advanced diaPASEF workflows report ∼3,400 proteins from pools of 1,000 cells.^7^ Concurrently with our study, a nanoLC-MS-based work recently reported ∼1,100 proteins from tumor-associated neutrophils with ∼30-min analysis time,^1^ indicating an emerging clinical applications.

Reducing separation time is an obvious route to higher throughput for routine applications. Current nanoLC methods quantify 1,000–5,000 proteins from single-cell–equivalent digests in 10–120 min gradients,^8–11^ while barcoded approaches achieve high cell counts—e.g., ∼1,000 proteins across 420 cells^12^ or ∼3,000 proteins across 1,490 cells^13^— typically at the cost of >1.5 h per cell in total runtime^12, 14–17^. Strategies such as dual-column LC^18^, short-gradient FAIMS-Orbitrap^19^, and nanowell or chip-based platforms^1, 19–22^ have pushed analysis to the 7.5–30 min range, establishing an aspirational benchmark of >1,000 proteins per cell at 100 cells/day.

Capillary electrophoresis (CE) offers a complementary route to speed. CE delivers high-efficiency separations with <30 s capillary conditioning, avoiding the relatively lengthy equilibrations required for LC. Our group and others have shortened CE-MS runs from ∼45 min to as little as 2 min, detecting ∼150 proteins from single-cell-equivalent inputs.^23–25^ Coupling CE to data-independent acquisition (DIA) has raised coverage to ∼1,250 proteins in 13–48 min.^23^ More recently, we introduced electrophoresis-correlative acquisition with artificial intelligence (Eco–AI), achieving 1,799 protein identifications from a single HeLa-cell-equivalent proteome digest in a 15-min CE run.^26^ Enabling this sensitivity with a∼7.2-min total runtime would theoretically support a 200 samples/day milestone in analytical throughput. Despite these advances, CE–MS has yet to unite the speed and sensitivity required for routine single-cell analysis (reviewed in ^9–11^).

Here, we report **Rapid Eco–AI**, a CE–MS platform tailored to speed for high-throughput and sensitivity for deep proteomic analysis of protein-limited specimens, single mammalian cells. Building on our earlier Eco–AI acquisition strategy,^26^ our goal was to achieve >1,250 protein identifications per cell at a CE–MS throughput advanced to ∼100 cells/day with single-plug to ∼200 cells/day using multi-plug samle injection. Further, we designed a hybrid proteome strategy to validate quantitative accuracy, before deploying the platform to freshly isolated human neutrophils from a healthy donor. To test the sensitivity of CE–MS, we intentionally profiled only ∼3 % of each cell’s proteome, the equivalent of sub-neutrophil proteomics (vs whole cells lately^1^). Collectively, these results establish Rapid Eco–AI CE–MS as a viable complement to nanoLC-MS for the emerging field of high-throughput single-cell proteomics, as exemplified here through the lens of immune cell heterogeneity.

## RESULTS AND DISCUSSION

### Development–Validation of a Rapid Eco–AI CE–MS Workflow

To achieve high-throughput, deep proteome coverage within a 5–7 min electrophoretic window, we adapted our previously reported 15-min Eco–AI method (**Fig. 1**). Rapid Eco–AI was trained on 300 pg of HeLa proteome digest—approximately the content of a single mammalian cell— leveraging real-time data-dependent acquisition (DDA) to dynamically prioritize MS² selection and AI-guided data processing for maximal proteome depth. Freshly isolated human neutrophils were individually isolated using an automated cell sorter (CellenOne, **Methods**). The cells were arrayed into precision-machined stainless-steel microwells, where they were shortly lysed and digested with trypsin *in situ* using nanoliter-level reagent volumes to minimize analyte loss. The resulting peptides were separated by CE, ionized via a cone-jet nanoESI interface, and analyzed on a Q Exactive Plus Orbitrap (Thermo Scientific) mass spectrometer, selected for its high resolving power and efficiency in CE–ESI coupling.^27^ Only ∼3% (∼2 pg) of each neutrophil’s proteome was measured in a single run, approximating to subcellular sampling, while profiling proteome heterogeneity among n = 13 neutrophils from a healthy volunteer.

**Figure 1.**
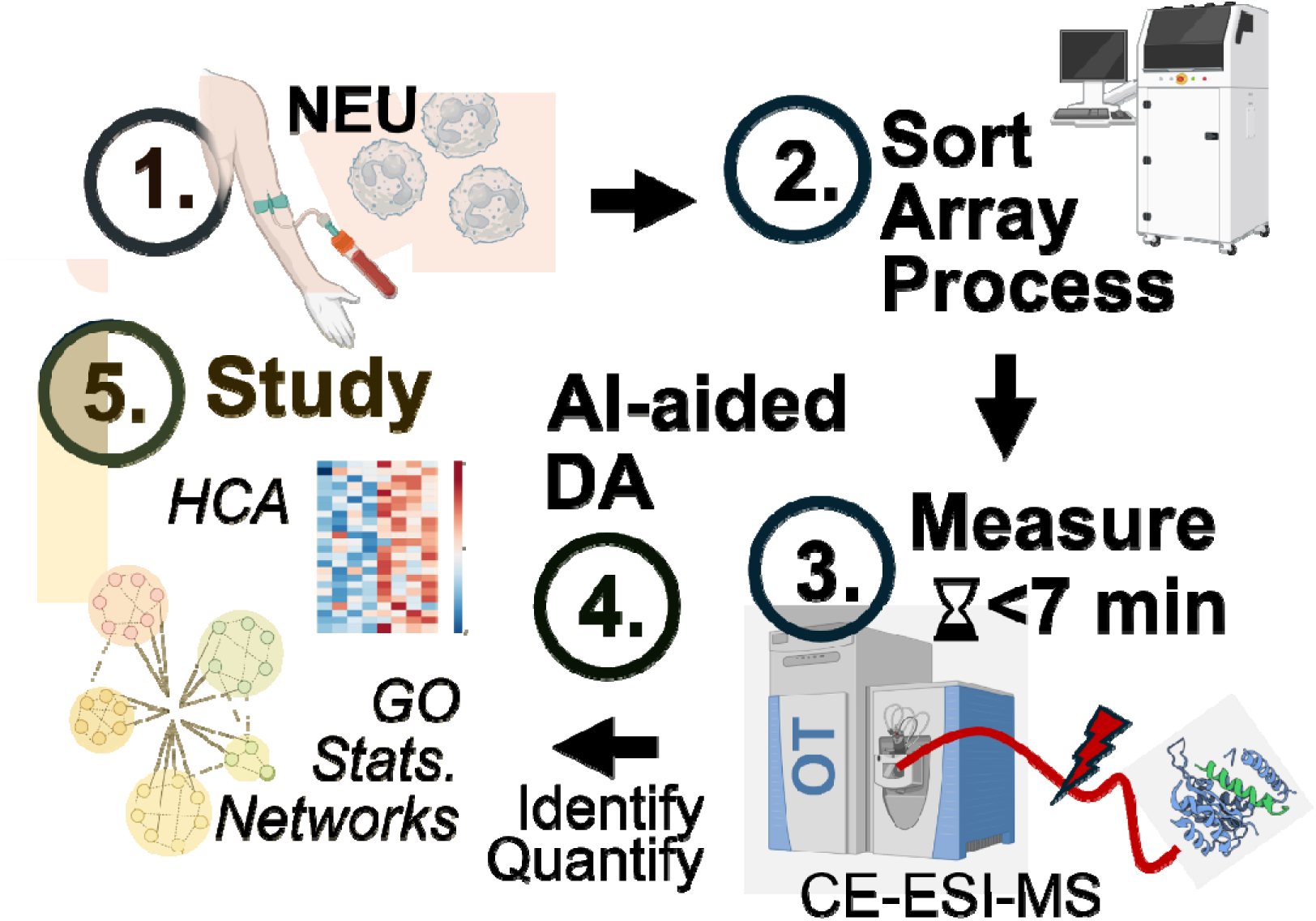
Workflow for proteomic profiling of human neutrophils using Rapid Eco–AI. Experimental steps are illustrated. (1) Fresh neutrophils (NEU) were isolated from whole blood and individually sorted into precision-machined stainless-steel microwells. (2) Each cell was trypsin-digested in nanoliter volumes for bottom-up proteomics. (3) Peptides were electrophoresed within a ∼7-min effective window in a fused silica capillary, ionized by electrospray (ESI), and analyzed on a Q Exactive Plus Orbitrap (OT) mass spectrometer (MS). Real-time data-dependent acquisition (DDA) guided MS² selection during Rapid Eco– AI, trained on 300 pg of HeLa digest. This achieved ∼1,350 protein identifications from single-cell–equivalent HeLa digests and ∼150 proteins from ∼2 pg (∼3 %) of a single neutrophil proteome. (4) AI-assisted data deconvolution (CHIMERYS) and label-free quantification (LFQ, DIA–NN) generated quantitative proteome profiles. (5) Chemometrics and pathway analyses revealed proteomic cell-to-cell heterogeneity with putative functional implications.

CE was fine-tuned to harmonize speed with sensitivity. Three electric field strengths—227, 357, and 429 V/cm—yielded effective separations of 15 min (<30 min total runtime), 7 min (<15 min), and 5 min (<11 min), respectively. In all cases, CE–ESI produced orderly migration patterns with precursor mass-to-charge (m/z) values increasing monotonically over migration time (MT) for each charge state (**Fig. 2**, +2 and +3 plotted). Slope analysis of these m/z–MT trends revealed increases of ∼84, 176, and 256 Th/min, corresponding to precursor shifts of 1.5, 3.0, and 4.3 Th/s. We reasoned that a 4 Th quadrupole isolation window, which we previously optimized^23, 26^, was sufficient to track these trend lines in real time, even under the most compressed separations.

**Figure 2.**
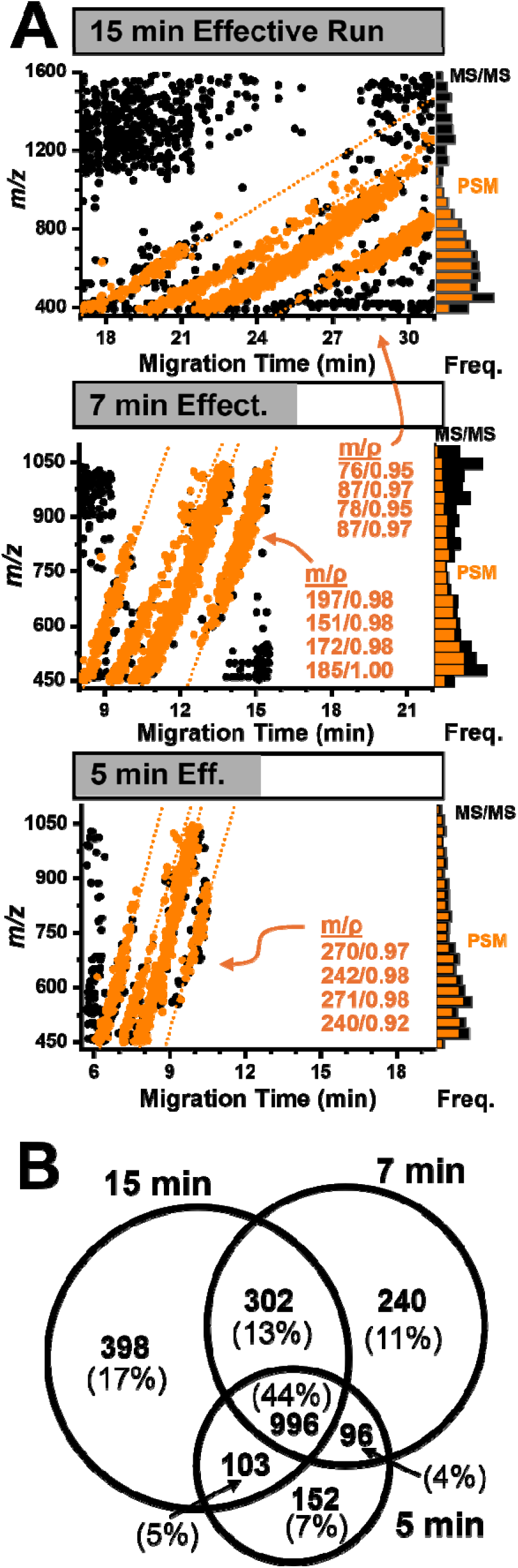
Refining data acquisition to Rapid Eco–AI. A constant proteome input (300 pg, ∼1 HeLa cell equivalent) was analyzed under varying electrophoretic conditions to achieve effective separation times of 15 min (25 kV, 110 cm capillary), 7 min (25 kV, 70 cm), and 5 min (30 kV, 70 cm capillary). **(A)** Analysis of the evolving precursor ion mass-to-charge (m/z) vs migration time (MT) data reflecting more compressed yet still highly ordered, charge-dependent Eco trends (**left plots,** charges labeled). Despite shortened runtimes, Rapid Eco– AI maintained efficient real-time precursor tracking and selection, yielding peptide spectral match (PSM) to MS² conversion rates between 62–81% between the peptide-rich *m/z* 400– 1,050 range (**right plots**). **(B)** Overlap among the proteins identified among technical triplicates, revealing robust identification with a high ∼45% overlap.

The experiments supported this notion. Narrowing the MS¹ survey range from *m/z* 350–1,600 to 400–1,050 enriched for peptide ions, improving the peptide spectral match (PSM)-to-MS² conversion from 62% to 81%. Importantly, the steeper m/z–MT slopes at 7 min and 5 min maintained high tracking efficiencies over separation time (**insets**; 70–80%), supporting robust and dynamic precursor ion sampling. **Figure 2B** presents the overlap among the identified proteomes. A high number of proteins were mutually detected (∼44% oerlap in proteome coverage), with a consistent, relatively low number of unique identifications (∼7– 17% coverage) between the 15-, 7-, and 5-min runs (**Table S1**). Combined, these metrics indicated robust performance despite the compressed separations.

Last, we iteratively refined MS detection to limited HeLa proteome inputs under 7 min of effective separation (**Fig. 3A**). The orbitrap analyzer was operated at different acquisition rates by combining Top-N selection (5 or 10) with MS² resolution (70,000 or 140,000 FWHM): Top-5/140K (slow, 3.1 s cycle), Top-10/70K (moderate, 2.8 s), and Top-5/70K (fast, 1.5 s). For 3 ng (∼10 cells) and 300 pg (∼1 cell) proteome inputs, the moderate and fast conditions outperformed the slow, yielding up to 1,828 and 1,634 cumulative protein identifications, respectively (**Table S2**). Sensitivity scaled well to smaller inputs, returning ∼1,357 proteins from 150 pg, or a half a cell and 835 proteins from ∼75 pg, or a quarter of a cell (**Table S2**)––the latter approximates to the total content of a single neutrophil for our demonstration. A 4 Th quadrupole isolation window provided the optimal balance between proteome coverage and cycle time (**Fig. 3B**), supporting our working hypothesis on efficient m/z–MT tracking using real-time DDA.

**Figure 3.**
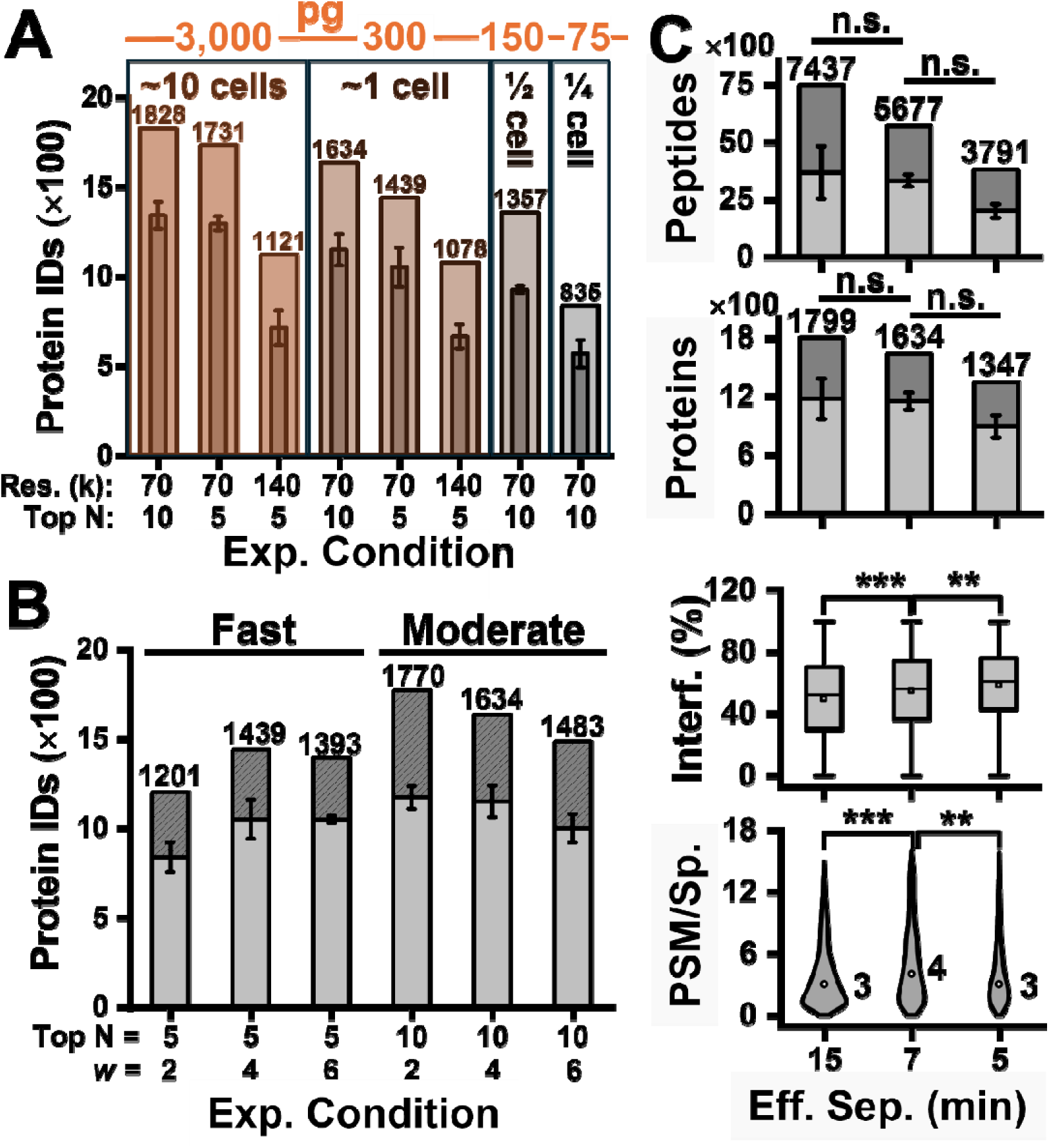
Optimization of MS acquisition parameters for Rapid Eco–AI. **(A)** HeLa proteome digests, from single-cell–equivalent (∼300 pg) down to subcellular amounts, were analyzed in triplicates on a Q Exactive Plus Orbitrap under three acquisition modes: fast (Top-5 at 70,000 FWHM; 1.5 s cycle), moderate (Top-10 at 70,000 FWHM; 2.8 s), and slow (Top-5 at 140,000 FWHM; 3.1 s). Fast and moderate settings yielded comparable protein identifications, with sensitivity scalable to the subcellular regime. **(B)** Effect of quadrupole isolation window (*w*) size on identification depth across analyzer speeds. **(C)** Peptide and protein identifications remained robust across 15-, 7-, and 5-min effective separations (**Table S1**), with high peptide spectral match (PSM)-to-MS² success rates despite slightly increasing MS^2^ interference (∼50%).

**Figure 4.**
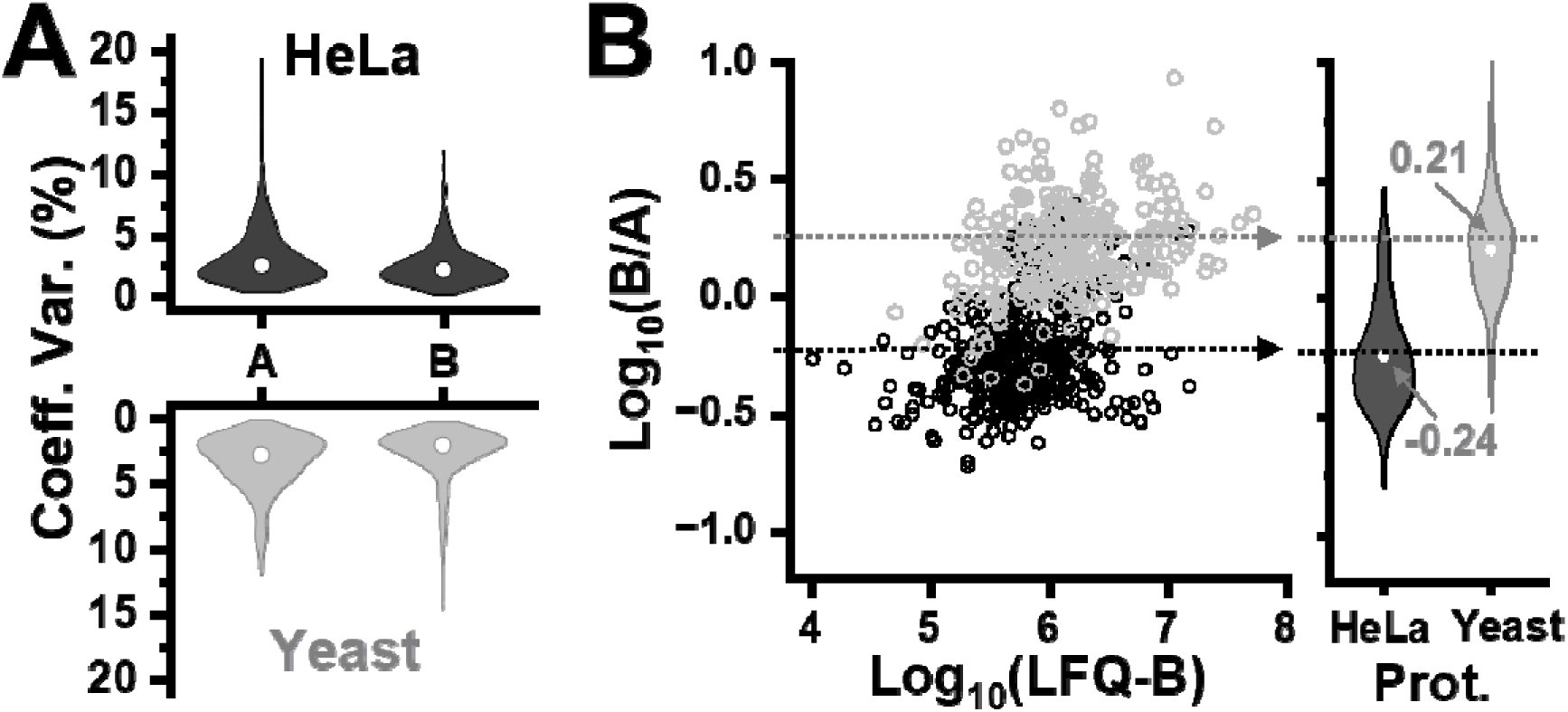
Validation of quantitative accuracy using a hybrid proteome model. **(A)** Percentage coefficient of variation (CV%) for peptides in samples containing HeLa and Yeas at “A” and “B” ratios across n = 5 technical replicates. For both HeLa and Yeast peptides, CVs were <20 % for all peptides, with median values of ∼2% after normalization. **(B)** Measured B/A peptide abundance ratios agreed closely with the theoretical values, validating accuracy. Key: dashed lines, expected ratios of 0.6:1 for human peptides and 1.8:1 for yeast peptides.

Despite the short separations, AI-guided data analysis remained robust (**Fig. 3C**). Protein and peptide identifications were comparable between the 15- and 7-min runs, with up to 7,437 peptides assigned to ∼1,800 proteins cumulatively (**Table S1**). Even at a separation lasting only ∼5 min, Rapid Eco–AI identified ∼1,347 proteins from 3,791 peptide sequences. The CHIMERYS AI-assisted algorithm (**Methods**) proved to tolerate increasing spectral interferences (∼50%) during searching increasingly chimeric tandem mass spectra manifesting from from faster separations. Although most MS^2^ spectra contained ca. <6 peptide spectral matches (PSMs), Rapid Eco-AI enabled the detection of up to 15 different peptides.^27^

Based on detected molecular features—m/z values with unique MT extracted in MaxQuant³ —most peptides produced transient signals lasting ∼12 s. Thus, our orbitrap cycles samples this signal with 4 (fast) to 8 (slow) discrete sampling events per feature.

Satisfying Nyquist–Shannon’s sampling theorem supported the reproducibility of label-free quantification (LFQ, DIA-NN, **Methods**). Indeed, the coefficient of variation (CV) between repeated LFQ of ∼300 pg HeLa proteome digest had a distribution mean at ∼17%, which standard data normalization improved to ∼2.3% (**Methods**, **Fig. S1**).

Further, we validated the accuracy of quantification using a hybrid HeLa:Yeast dual-proteome model. Known amounts of HeLa and Yeast proteome digests^28–29^ were mixed to prepare samples “A” and “B” with expected B/A peptide abundance ratios of 0.6 for human and 1.8 for yeast. Each sample was repeatedly analyzed under 7 min effective electrophoresis in 5 technical replicates (**SI Methods**). Peptide abundances were estimated by label-free quantification (LFQ) using CHIMERYS (**Table S3**). Following log transformation, median coefficients of variation (CVs) were <5 %. The measured log B/A ratios closely matched the expected values—0.64 vs. 0.60 for human peptides and 1.78 vs. 1.80 for yeast. Together, reproducible and accurate quantification with deep sensitivity and speed align with benchmarks recommended for robust and rigorous science by a recent single-cell proteomics working group^30^.

### Human Neutrophil Profiling

To demonstrate proof-of-principle, we performed proteome profiling of individual human neutrophils. The cells, 16–23 µm in diameter (**Fig. S4**), were isolated from a healthy donor and deposited into stainless-steel microwells using an automated cell sorter (cellenONE, **Methods**; **Fig. 5A**). Each cell was digested *in situ* with 250 nL of 1 ng/µL trypsin co-deposited in automation; to prevent evaporation of the digestion media over 1 h of incubation, 1 µL of 50 mM TEAB was added to each well. On completion, the cellular proteome digests were air dried, reconstituted in 500 nL of CE–MS solvent^26, 31^, and a 15 nL aliquot (∼2 pg, ∼3 % of each cell’s proteome) was analyzed over a 7-min electrophoretic window.

**Figure 5.**
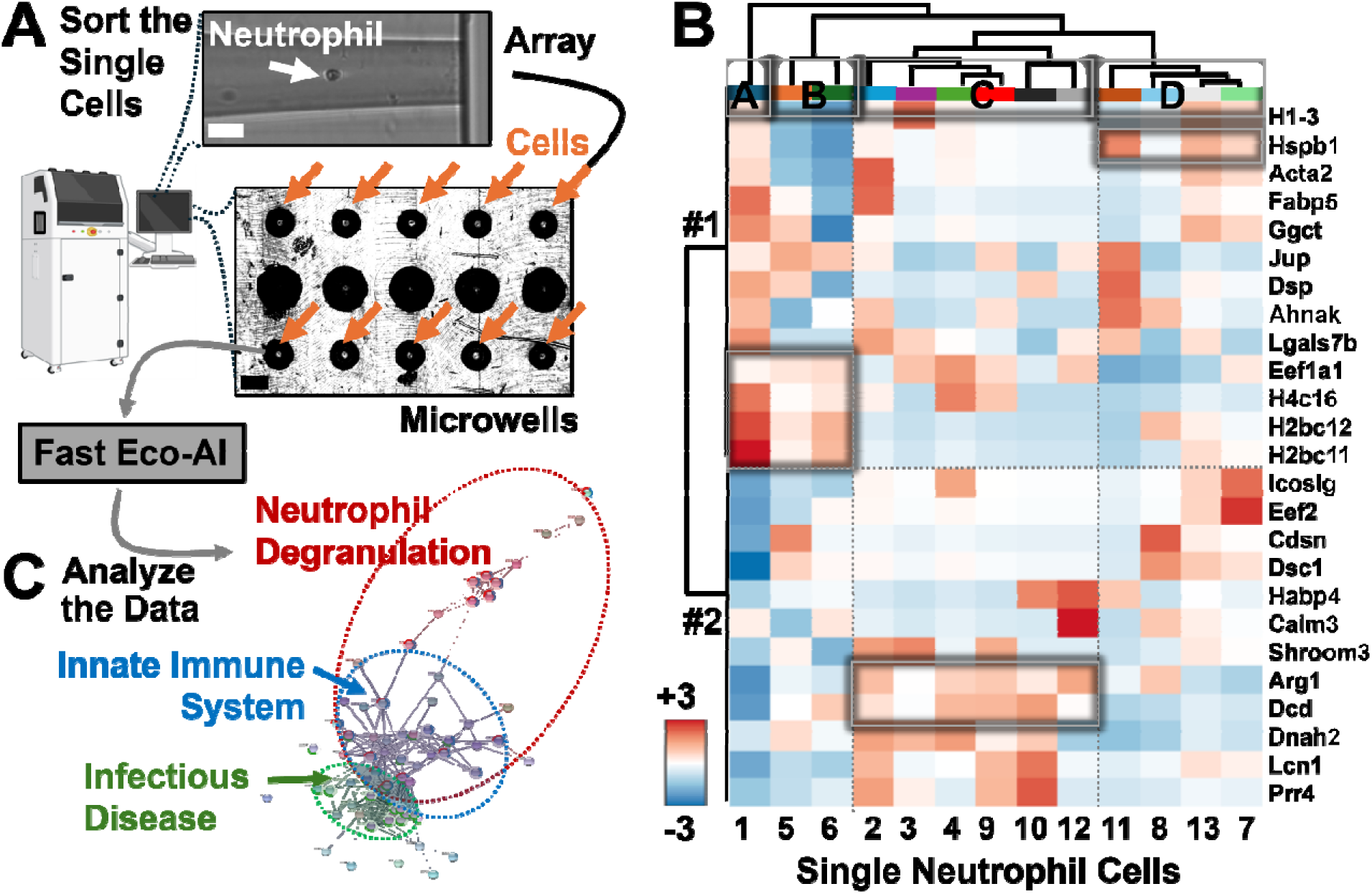
Single-cell proteomic profiling of human neutrophils using Rapid Eco–AI. **(A)** Freshly isolated neutrophils from a healthy volunteer were individually arrayed into micromachined wells using an automated cell sorter (cellenONE). The upper inset shows a representative neutrophil in the sorting capillary before deposition; the lower inset highlights microwells containing single cells (arrows). A total of *n* = 13 neutrophils were analyzed by CE–MS with a 7-min effective CE window. Scale bars: 40 µm (white), 1 mm (black). **(B)** Unsupervised hierarchical cluster (HCA) and heatmap analysis of the top 25 most variable proteins revealed at least four distinct neutrophil subgroups (labeled A–D), supported by differential protein expression patterns (regions #1–2), including histones (H4c16, H2bc11/12), HSPB1, DCD, and ARG1. **(C)** STRING network analysis of known/predicted protein-protein associations among the 132 identified proteins revealed enrichment in canonical neutrophil pathways, including degranulation, innate immune response, and infectious disease processes.

Comparison of peptide m/z distributions between HeLa and neutrophil digests demonstrated that CE–MS parameters optimized on HeLa were reasonably transferable to neutrophil analysis (**Fig. S3**). Indeed, across n = 13 single neutrophils, Rapid Eco–AI identified 151 proteins (**Table S4**), with per-cell IDs ranging from 5 to 79. Interspersed blank injections confirmed the absence of carryover between consecutive analyses. The observed proteome depth variability tracked with cell-size heterogeneity (16–23 µm diameter; **Fig. S4**), reflecting up to a 3-fold difference in total protein content and likely contributing to proteome coverage differences.

To interrogate the biological significance of the observed proteomic profiles, label-free quantification (LFQ) intensities were median-normalized, log-transformed, and autoscaled. Unsupervised hierarchical clustering (HCA) of the 25 most variable proteins (**Fig. 5B**) resolved the cells into 4 groups (A–D), with protein clusters #1–2 defining each subgroup. Groups A and B were enriched in histone proteins—Histone 4 (H4C16) and H2B variants (H2BC11/12), region #1. In contrast, Group D (region #2) exhibited higher HSPB1 (HSP27) abundance, a chaperone implicated in regulating neutrophil chemotaxis.^32^ Group C (region #3) was characterized by elevated dermcidin (Dcd), an antimicrobial peptide in innate immunity,^33^ along with increased arginase 1 (Arg1), which attenuates T-cell responses via L-arginine hydrolysis.^34^ Together, these data appear to suggest distinct functional states and basal heterogeneity among the individual neutrophils. Importantly, subgroups A/B and C classifications is consistent with a recent report describing neutrophil populations based on the same protein markers: increased histones and Arginase 1, respectively.^1^

STRING^35^ analysis of the combined dataset revealed significant enrichment in neutrophil degranulation, innate immunity, and infectious disease pathways (**Fig. 5C**, **Table 1**). Among the 151 identified proteins, several key mediators of neutrophil activation were detected.

**Table 1.** STRING enrichment analysis of canonical protein-protein interactions on the measured neutrophil cells.

| Cellular Component | FDR | Gene Counts |
| --- | --- | --- |
| Ficolin-1-rich granule | $3.51 \times 10^{-8}$ | 14 |
| Secretory granule lumen | $1.16 \times 10^{-8}$ | 18 |
| Azurophil granule lumen | $3.92 \times 10^{-6}$ | 9 |
| Vacuolar lumen | $1.30 \times 10^{-6}$ | 12 |
| Secretory granule | $6.68 \times 10^{-8}$ | 27 |

| Reactome Pathways | FDR | Gene Counts |
| --- | --- | --- |
| Neutrophil degranulation | $1.57 \times 10^{-11}$ | 26 |
| Innate Immune System | $5.58 \times 10^{-9}$ | 32 |
| Influenza Infection | $8.39 \times 10^{-9}$ | 14 |
| Infectious disease | $1.44 \times 10^{-8}$ | 29 |
| Cellular responses to stress | $1.74 \times 10^{-6}$ | 26 |
| Immune System | $1.24 \times 10^{-5}$ | 37 |

Macrophage migration inhibitory factor (Mif) – a cytokine that recruits neutrophils and amplifies inflammation – was consistently observed.^36^ Members of the S100 calcium-binding family were also prominent, including psoriasis (S100A7), which promotes neutrophil extracellular trap (NET) formation,^37^ calgranulin B (S100A9), a canonical activation marker,^37^ and calcyclin (S100A6), involved in neutrophil recruitment and polarization.^38^ Additionally, peroxiredoxin 6 (PRDX6), known to translocate to the plasma membrane during activation to modulate NADPH oxidase activity,^39^ was identified across multiple cells. These findings demonstrate that Rapid Eco–AI provides sufficient depth and precision to capture critical functional proteins in single neutrophils, enabling future studies of immune-cell phenotypes at single-cell resolution.

## CONCLUSIONS

We have developed **Rapid Eco–AI**, a fast CE–MS workflow that delivers robust, high-sensitivity single-cell proteomics in under 7 min of effective separation. We found Real-Time Eco-DDA sufficiently dynamic to boost CE-ESI sensitivity during data acquisition. With AI-driven data deconvolution, Eco–AI consistently identified and quantified ∼1,350 proteins from ∼300 pg of HeLa digest with median CVs <20%. Sensitivity scaled to ∼835 proteins from ∼75 pg of input, demonstrating compatibility with small, single-cell–level proteome amounts, such as those contained in single human neutrophils.

Applying Rapid Eco–AI to 13 individual human neutrophils yielded the first CE–MS–based single-cell proteome profiles of this leukocyte. The analysis revealed a conserved core of immune-related proteins alongside marked intercellular variability. Key activation markers— S100 family members, HSPB1, and ARG1—were differentially expressed across 4 subpopulations defined by unsupervised chemometrics. These results highlight functional heterogeneity within a morphologically uniform neutrophil population and underscore the need for further studies to elucidate the biological roles of these sub-phenotypes.

As demonstrated here, Rapid Eco–AI is a scalable, sensitive, and rapid platform for both single-cell and subcellular proteomics, affordably^27^. Its performance can be further enhanced through automated nanoliter-scale sample handling (e.g., nanoPOTS^40^, SCopE MS^17, 41^), advanced microanalytical CE systems (e.g., RoboCap^42^), and coupling to contemporary mass analyzers (time-of-flight^16, 43^ and Orbitrap^44^) with higher speed and dynamic range.

Integration of ion-mobility separation (e.g., FAIMS^45–46^) or project-specific spectral libraries^47^ will simplify chimeric spectra and deepen coverage. We anticipate that Rapid Eco– AI will be broadly adaptable to diverse cell types, enabling high-throughput functional immunophenotyping and advancing applications in systems immunology, inflammation biology, and clinical proteomics.

## ACKNOWLEDGMENT

Parts of this research were supported by the University of Maryland (seed funding to P.N.) or the National Institute of Aging (R01AG088147 to P.N.) or the National Institute of General Medical Sciences award no. (R35GM124755 to P.N.) of the US National Institutes of Health or Fulbright Brazil (Fellowship to I.L., host P.N.) or Distrito Federal Research Foundation (FAPDF, to W.F.). The conclusions are the authors’ responsibility and do not necessarily reflect those of the sponsors. We thank Justin Parks, Sara Mascone, Joshua Cantlon, and Sushant Ranadive for assistance during sample collection.

---

**TOC Image Description:**
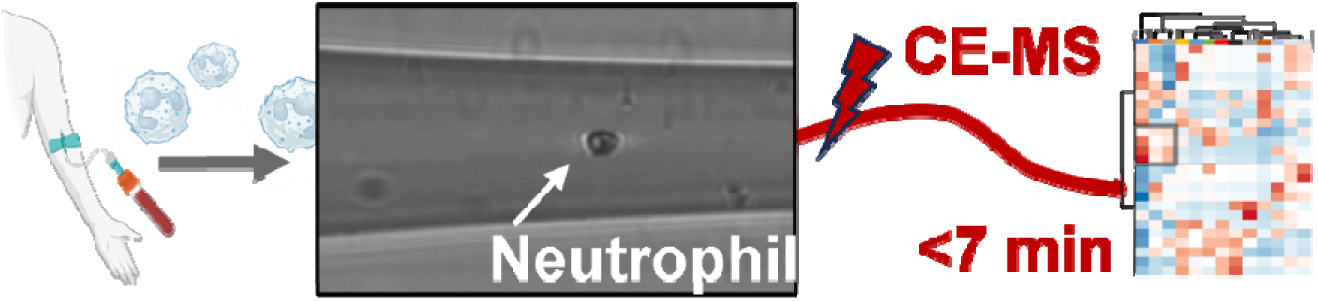
Rapid capillary electrophoresis driving electrophoresis-correlative mass spectrometry profiles human neutrophils, revealing distinct proteomic subpopulations in <7-min effective analyses.

