## Supplementary material for "Rapid CE–MS with Real-Time Eco–AI Resolves Proteomic Heterogeneity Among Single Human Neutrophils": SI Document

### SI METHODS

**Materials.** EDTA tubes were purchased from BD Vacutainer (catalog no. 366643, BD Vacutainer, Franklin Lakes, NJ). Lymphocyte Separation Medium (LSM) was obtained from Corning (cat. no. 25-072-CV, Corning, Manassas, VA). The Fast Panoptic Staining Kit for Haematology was acquired from Glentham Life Sciences (cat. no. GX5533, Corsham, UK). Trypsin Platinum was purchased from Promega (Madison, WI, USA). Dextran (MW ca. 500,000 g/mol) and 0.4% Trypan Blue solution (cat. no. 15250061, Waltham, MA) were used as indicated. High-purity solvents suitable for HPLC-MS were sourced from Thermo Fisher Scientific: acetic acid (AcOH), acetonitrile (ACN), formic acid (FA), methanol (MeOH), and water. The HeLa proteome digest standard (cat. no. 88329, Pierce, Rockford, IL) was from Thermo Fisher, and the Yeast proteome standard (cat. no. V7461) was from Promega.

A Neubauer chamber used for counting polymorphonuclear leukocytes (PMNs) was supplied by JoanLab (Amazon, USA). Neutrophils were sorted using a cellenONE automated cell arrayer (Cellenion, Lyon, France) equipped with a medium-sized uncoated piezoelectric dispensing capillary (PDC). All solutions used during PMN isolation from whole blood were filtered through a sterile 0.02 μm filter (Semmerfeld, Amazon, USA).

Fused silica capillaries for capillary electrophoresis were obtained from Polymicro Technologies (40/105 μm inner/outer diameter, catalog no. 1068150596, Phoenix, AZ). Borosilicate glass capillaries (0.75/1.00 mm inner/outer diameter, catalog no. B100-75-10, Sutter Instrument, Novato, CA) were used to fabricate CE-nanoESI emitters.

**Solutions.** The erythrocyte sedimentation solution was composed of 0.15 M NaCl and 6% dextran (w/v). For PMN suspension and erythrocyte lysis, Hank's Balanced Salt Solution (HBSS) 1X and 2X were prepared from HBSS 10X (catalog no. H4641, Sigma-Aldrich, St. Louis, MO). A 0.2% trypan blue solution was prepared in 1× HBSS and used to assess cell viability.

The *background electrolyte (BGE)* used in capillary electrophoresis was composed of 1 M FA in 25% (v/v) ACN. For the CE-ESI interface, *the sheath solution* consisted of 0.5% (v/v) AcOH in 10% (v/v) MeOH. Both HeLa proteome digests and single-cell samples were reconstituted in a *sample solvent* containing 0.05% (v/v) FA in 75% (v/v) ACN.

Venous blood was collected from a healthy volunteer into a tube containing dipotassium EDTA, following protocols approved by the Institutional Review Board (approval no. 2124276) and the Institutional Biosafety Committee (project no. PN645) of the University of Maryland, College Park. The blood sample was centrifuged at 1,500 × g for 15 min. The resulting buffy coat was transferred to a clean conical tube containing 5 mL of phosphate-buffered saline (PBS) and gently mixed by inversion (4–5 times). The diluted sample was carefully layered over 3 mL of Lymphocyte Separation Medium (LSM) and centrifuged at 1,400 × g for 15 min. The clear upper phases, containing plasma and lymphocytes, were reserved for other studies. The lower 2 mL of the dark phase, containing red blood cells (RBCs) and polymorphonuclear leukocytes (PMNs), was transferred to a tube containing 0.15 M NaCl and 6% dextran (cat. no. J63702.09, MW ca. 500,000 g/mol, Thermo Scientific Chemicals, Waltham, MA) for RBC sedimentation at room temperature over 30 min. The clear upper phase containing PMNs was processed as previously described.[1] Briefly, the cell pellet was washed twice with 1× HBSS, and residual RBCs were removed by hypotonic lysis. A 3 μL aliquot of the cell suspension was placed on a clean glass slide and stained using the Fast Panoptic Staining Kit for Haematology (cat. no. GX5533,

Glentham Life Sciences, Corsham, UK) for evaluation of morphology and purity by light microscopy. For cell viability and yield assessment, 5 µL of the PMN suspension was mixed with 45 µL of 0.2% trypan blue solution (cat. no. 15250061, Thermo Fisher Scientific, Waltham, MA) and analyzed in a Neubauer chamber (JoanLab, Amazon, USA) under light microscopy. The sample was deemed suitable for further analysis if it exhibited ≥95% cell viability and neutrophil purity. The cell suspension was then washed twice with PBS, and the concentration was adjusted to 150 cells/µL prior to single-cell sorting.

**Single Neutrophil Sorting and Processing.** Single neutrophil sorting, isolation, and proteomic processing were performed using the cellenONE system (Scienion US Inc., Tempe, AZ). The instrument chamber was maintained at 65% relative humidity. A medium-sized, uncoated piezoelectric dispensing capillary (PDC) was used under the following conditions: voltage, 148 V; pulse length, 49 µs; frequency, 500 Hz; LED delay, 200 µs; and LED pulse width, 4 µs. Cells were selected using the transmission imaging channel with inclusion criteria of 14–23.1 µm diameter and elongation values below 2.50 (maximum threshold = 7). Single cells were dispensed into a custom-fabricated 30-well processing plate, with 21 wells loaded per run. Each well received 250 nL of degassed trypsin solution (1 ng/µL in 50 mM TEAB). Following digestion, 1 µL of 50 mM TEAB was added to each well to prevent evaporation. The plate was sealed with parafilm and incubated at 40 °C for 1 h. After digestion, the neutrophil proteomes were stored at –80 °C until CE-MS analysis.

**Hybrid Proteome Preparation.** To validate the accuracy of label-free quantification using maxLFQ approach, a HeLa–Yeast dual-proteome digest sample was prepared at known ratios. To obtain HeLa:Yeast proteome reference, 8 µg of the HeLa digest was spiked with 4 µg of the yeast to produce sample "A" or 12 µg for sample "B". The reference was diluted in the CE sample buffer containing a final peptide concentration of 0.3 µg/µL for CE-MS analysis. Therefore, upon analysis, the expected ratios B/A are 0.6 for HeLa and 1.8 for yeast.

**CE-ESI-MS Analysis.** The proteome digests from single neutrophils, the HeLa standard, and the hybrid proteome were reconstituted in the *sample solvent* and then analyzed on the same custom CE-nanoESI platform, following protocols previously described.[2, 3] The HeLa proteome digests (75 pg–3 ng) and the hybrid proteome (3 ng) were separated in a BGE-filled 75-cm fused silica capillary at an electric field strength of 400 V/cm. The capillary outlet was interfaced with a sheath-flow nanoESI emitter powered by an electrokinetic pump (+200–500 V) and operated in the cone-jet spraying regime for efficient ionization, based on previously described designs.[4, 5] The electrospray Taylor cone was monitored in real time using a long-working-distance objective (Mitutoyo Plan Apo) and CCD camera (EO-2018C, Edmund Optics). The emitter was positioned ~1 mm from the MS orifice using a 3-axis translation stage.

The ionized peptides were analyzed using a Q Exactive Plus hybrid quadrupole–orbitrap mass spectrometer (Thermo Fisher Scientific) operated in data-dependent acquisition (DDA) mode. The following Top-N configurations were tested based on complementary analytical speed (top-N/Orbitrap resolution): slow (***Top-5/140k),*** Top-5/140,000 FWHM yielding ~3.1 s cycle time; moderate speed (***Top-5/70k)***, Top-5/70,000 FWHM yielding ~2.8 s cycle time; and fast (***Top-10/70k)***, Top-10/70,000 FWHM yielding ~1.5 s cycle time. Quadrupole mass-to-charge (m/z) isolation widths of 2, 4, and 6 Th were tested. All other acquisition parameters were held constant: survey scan range ($MS^1$), *m/z* 400–1,050; $MS^1$ resolution, 70,000 FWHM; C-trap maximum injection time, 240 ms; $MS^2$ AGC target, $3 \times 10^6$ counts; precursor charge states, +2 to

+4; and higher-energy collisional dissociation (HCD) energy, 28% normalized collision energy (NCE).

**Data Analysis.** The MS data were searched against the HeLa proteome (UP000005640, UniProt, July 2023 release, 20,523 entries) or the yeast proteome (UP000002311, UniProt, April 2025 downloaded, 6,067 entries) using Proteome Discoverer 3.0 (Thermo Fisher Scientific), employing the CHIMERYS node (prediction model: *inferys_2.1_fragmentation*) for protein identification. The query parameters were: fixed modification, cysteine carbamidomethylation, variable oxidation, methionine, peptide length, 7–30 amino acids, missed tryptic cleavages, max 2, fragment ion mass tolerance, 20 ppm. Common contaminant detection was enabled. A minimum of one proteotypic peptide was required to identify a protein. All protein identifications were filtered to a false discovery rate (FDR) <1%, determined using a reversed-sequence decoy proteome database.

Label-free protein quantification was performed using LFQ in DIA-NN 1.832 in library-free mode with the following parameters: peptide length (5–35), maximum number of variable modifications (1), missed cleavages (2), and precursor charge range (+2 to +4). All other settings were set to the default.[6] The resulting label-free quantification (LFQ) abundance values were median-normalized to compare equal amounts of proteomes, $\log_{10}$-transformed to factor over broad concentration ranges, and auto-scaled in MetaboAnalyst 6.0.[7] Data visualization and statistical analyses were performed in OriginPro 2020b (Origin Lab, Northampton, MA). The Mann–Whitney U test was conducted in R, with exact $p$-values reported unless computational limits applied. A $p < 0.05$ was considered statistically significant in this project.

**Scientific Rigor.** HeLa and hybrid proteome digests were analyzed in 3–5 technical replicates (i.e., the same sample was measured multiple times). A total of $n = 13$ single neutrophils (biological replicates) were sorted and processed from freshly collected human blood from the same batch from the same healthy volunteer. Sample acquisition and measurement order were randomized. Both biological and technical replicates were randomized. The CE-ESI-MS platform was rinsed between each analysis. Analysis of the BGE alone (blanks) confirmed no detectable analyte carryover between consecutive runs. Common contaminants (CRAP) were removed and are excluded from the final dataset.

**Safety.** All biological materials were handled, processed, and deactivated following institutionally approved biosafety protocols. Chemicals were managed following standard laboratory safety practices. Special care was taken when handling capillaries and ESI emitters to minimize puncture risk. All electrically conductive components of the CE-MS platform were grounded or enclosed within safety-interlocked housings to prevent electrical hazards.

**Data Availability.** The MS files of the standard digests and neutrophil proteomes were deposited in the ProteomeXchange Consortium via the PRIDE partner repository under accession number identifier PXD067364.

### SI REFERENCES

1. Souza Luz, I.; Takaya, R.; Gonzaga Ribeiro, D.; Sales Silva, N.; Fontes, L.; Castro, M. S.; Fontes, W., A set of screening techniques for a quick overview of the neutrophil function. *JoVE* **2024,** (204), No. e65329.

### SI FIGURES

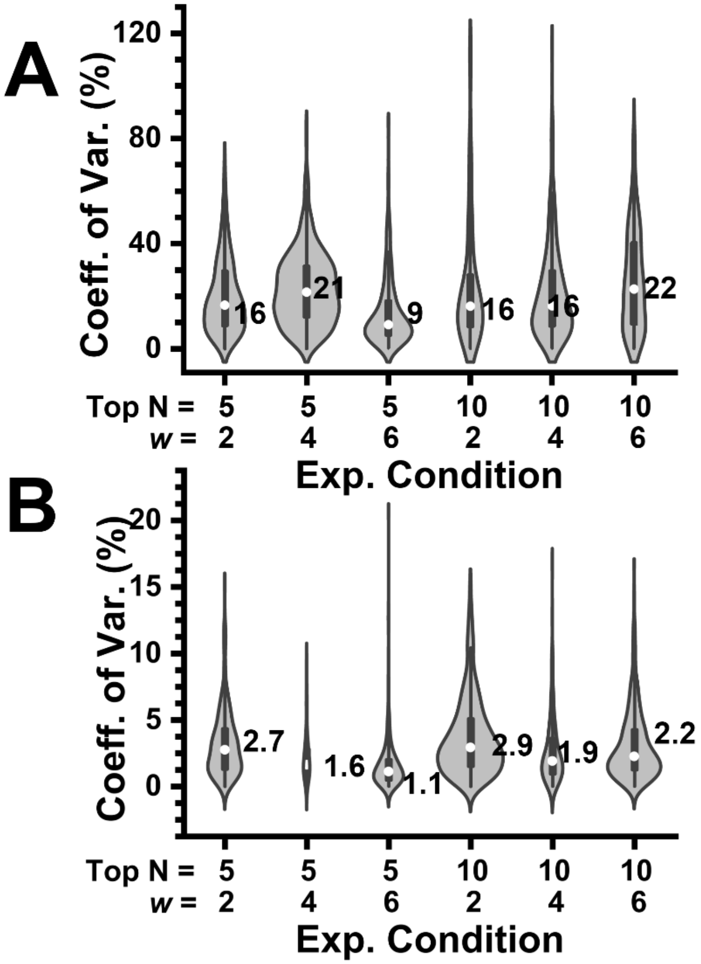

**Figure S1. Optimization of proteome reproducibility for Rapid Eco–AI.** Experimental parameters were optimized using single-cell–equivalent amounts of HeLa digest (~300 pg), each analyzed within a ~7-min effective electrophoretic window. Precursor abundance thresholds were tested using signal-based Top-N selection with quadrupole isolation widths of 2, 4, and 6 Th. **(A)** before and **(B)** after data normalization (median-normalized, $log_{10}$-transformed, and auto-scaled).

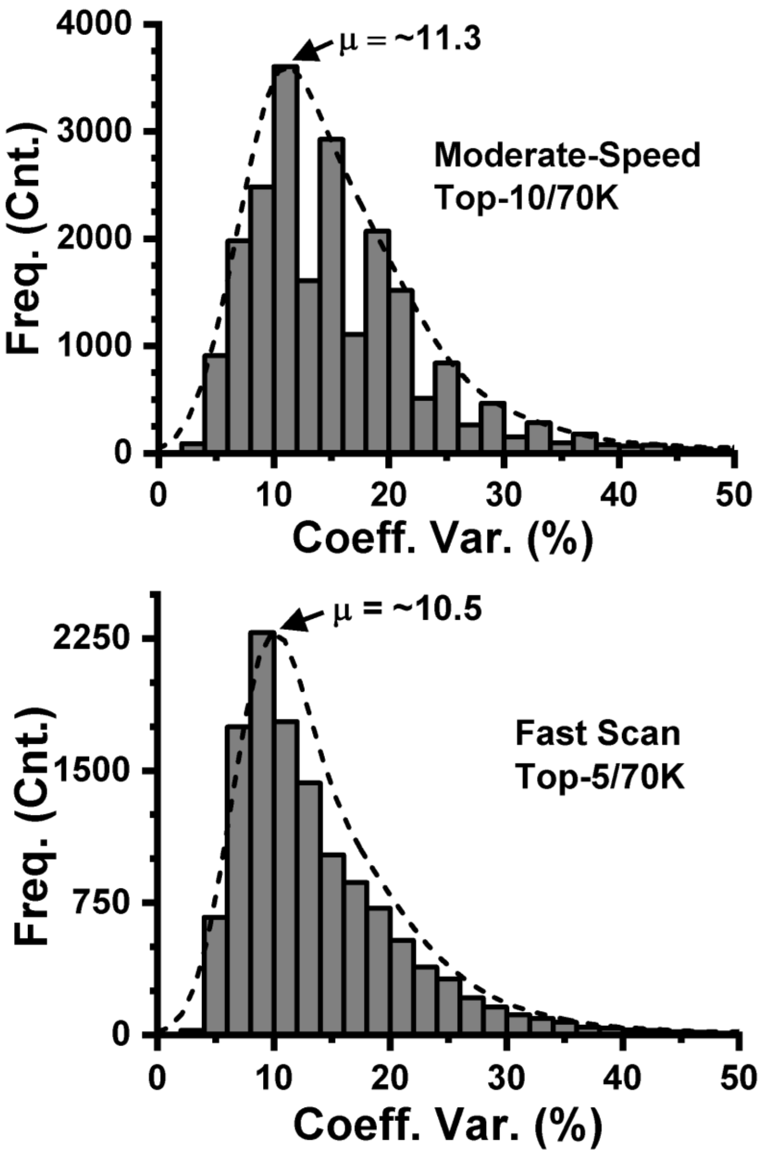

**Figure S2. Analysis of CE-MS peak width distribution.** With a mean electrophoretic peptide width of ~12-s, the orbitrap analyzer operating at moderate (Top-10/70K) and fast (Top-5/70k) speed acquired 4–7 $MS^2$ data points per peak. These results support the suitability of the Rapid Eco–AI method for accurate quantitation despite its short electrophoretic separation window.

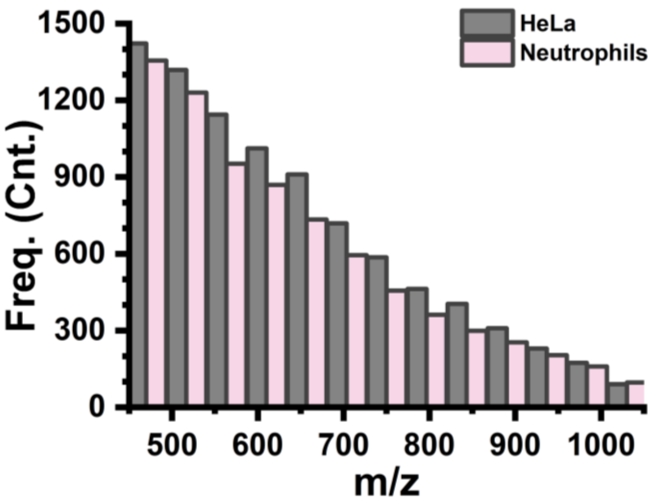

**Figure S3. Comparison of detected peptide m/z distributions for HeLa and human neutrophil proteome digests**. Similar mass-to-charge (m/z) distribution suggested the CE–MS parameters optimized on HeLa were transferable to neutrophil proteome analysis.

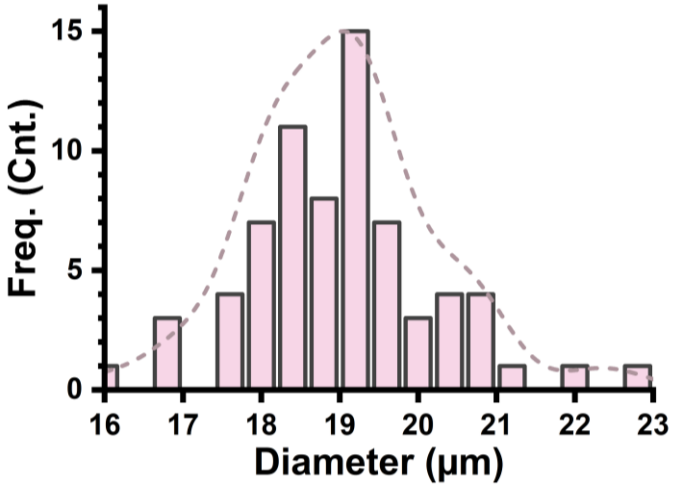

**Figure S4. Neutrophil size distribution.** A total of n = 70 different cells showed a diameter range of 16–23 µm with a median size of ~19.5 µm during CellenONE analysis.
